# DNNBrain: a unifying toolbox for mapping deep neural networks and brains

**DOI:** 10.1101/2020.07.05.188847

**Authors:** Xiayu Chen, Ming Zhou, Zhengxin Gong, Wei Xu, Xingyu Liu, Taicheng Huang, Zonglei Zhen, Jia Liu

## Abstract

Deep neural networks (DNNs) have attained human-level performance on dozens of challenging tasks through an end-to-end deep learning strategy. Deep learning gives rise to data representations with multiple levels of abstraction; however, it does not explicitly provide any insights into the internal operations of DNNs. Its success appeals to neuroscientists not only to apply DNNs to model biological neural systems, but also to adopt concepts and methods from cognitive neuroscience to understand the internal representations of DNNs. Although general deep learning frameworks such as PyTorch and TensorFlow could be used to allow such cross-disciplinary studies, the use of these frameworks typically requires high-level programming expertise and comprehensive mathematical knowledge. A toolbox specifically designed for cognitive neuroscientists to map DNNs and brains is urgently needed. Here, we present DNNBrain, a Python-based toolbox designed for exploring internal representations in both DNNs and the brain. By integrating DNN software packages and well-established brain imaging tools, DNNBrain provides application programming and command line interfaces for a variety of research scenarios, such as extracting DNN activation, probing DNN representations, mapping DNN representations onto the brain, and visualizing DNN representations. We expect that our toolbox will accelerate scientific research in applying DNNs to model biological neural systems and utilizing paradigms of cognitive neuroscience to unveil the black box of DNNs.

## Introduction

Over the past decade, artificial intelligence (AI) has made dramatical advances because of the rise of deep learning (DL) techniques. DL makes use of a deep neural network (DNN) to model complex non-linear relationships, and thus is able to solve real-life problems. A DNN often consists of an input layer, multiple hidden layers, and an output layer. Each layer generally implements some non-linear operations that transform the representation at one level into another representation at a more abstract level. As a result, DL could automatically discover multiple levels of representations that are needed for a given task (LeCun et al., 2015; Goodfellow et al., 2016). Particularly, the deep convolutional neural network (DCNN) architecture stacks multiple convolutional layers hierarchically, inspired by the hierarchical organization of the primate ventral visual stream. A supervised learning algorithm is generally used to tune the parameters of the network to minimize the error between the network output and the target label in an end-to-end manner (LeCun et al., 1998; Rawat and Wang, 2017). With such a built-in architecture and learning on large external datasets, DCNNs have achieved human-level performance on a variety of challenging object (Krizhevsky et al., 2012; Szegedy et al., 2014; Simonyan and Zisserman, 2015; He et al., 2016) and speech recognition tasks (Hinton et al., 2012; Sainath et al., 2013; Hannun et al., 2014).

Besides the feats of engineering, DNNs provide a potentially rich interaction between studies on biological and artificial information processing systems. On the one hand, DNNs offer the best models of biological intelligence so far (Cichy and Kaiser, 2019; Richards et al., 2019). Particularly, good correspondence has been identified between DNNs and the visual system (Yamins and DiCarlo, 2016; Kell and McDermott, 2019; Serre, 2019; Lindsay, 2020). First, DNNs exhibit similar behavioral patterns to human and non-human primate observers on some object recognition tasks (Jozwik et al., 2017; Rajalingham et al., 2018; King et al., 2019). Second, DCNNs appear to recapitulate the representation of visual information along the ventral stream. That is, early stages of the ventral visual stream (e.g., V1) are well predicted by early layers of DNNs optimized for visual object recognition, whereas intermediate stages (e.g., V4) are best predicted by intermediate layers, and late stages (e.g., IT) are best predicted by late layers (Khaligh-Razavi and Kriegeskorte, 2014; Yamins et al., 2014; Güçlü and van Gerven, 2015; Eickenberg et al., 2017). Finally, DNNs designated for object recognition spontaneously generate many well-known behavioral and neurophysiological signatures of cognitive phenomena such as shape tuning (Pospisil et al., 2018), numerosity (Nasr et al., 2019), and visual illusions (Watanabe et al., 2018), and thus provide a new perspective to study the origin of intelligence. Indeed, neuroscientists have already used DNNs to model the primate visual system (Schrimpf et al., 2018; Lindsey et al., 2019; Lotter et al., 2020).

On the other hand, the end-to-end DL strategy makes DNN a black box, without any explanation of its internal representations. The experimental paradigms and theoretical approaches from cognitive neuroscience have significantly advanced our understanding of how DNNs work (Hasson and Nusbaum, 2019). First, concepts and hypotheses from cognitive neuroscience, such as sparse coding and modularity, provide a hand-on terminology to describe the internal operations of a DNN (Agrawal et al., 2014; Ritter et al., 2017). Second, a variety of methods in manipulating stimuli such as stimulus degradation and simplification have been used to characterize units’ response dynamics (Baker et al., 2018; Geirhos et al., 2019). Finally, rich data analysis techniques in cognitive neuroscience, such as ablation analysis (Morcos et al., 2018; Zhou et al., 2018), activation maximization (Nguyen et al., 2016), and representation similarity analysis (Khaligh-Razavi and Kriegeskorte, 2014; Jozwik et al., 2017), provide a powerful arsenal in exploring the computational mechanisms of DNNs.

Such a crosstalk between cognitive neuroscience and AI needs an integrated toolbox that can fulfill the requirements of both fields. However, the most commonly-used DL frameworks such as PyTorch^1^ and TensorFlow^2^ are developed for AI researchers. The use of these frameworks typically requires high-level of programming expertise and comprehensive mathematical knowledge of DL. To our knowledge, there is no software package that is specifically designed for both AI scientists and cognitive neuroscientists to interrogate DNNs and brains at the same time. Therefore, it would be of great value to develop a unifying toolbox to integrate DNN software packages and well-established brain mapping tools.

Here we present DNNBrain, a Python-based toolbox specifically designed for exploring representations in both DNNs and brains (Figure 1), which has five major features.

**Figure. 1.**
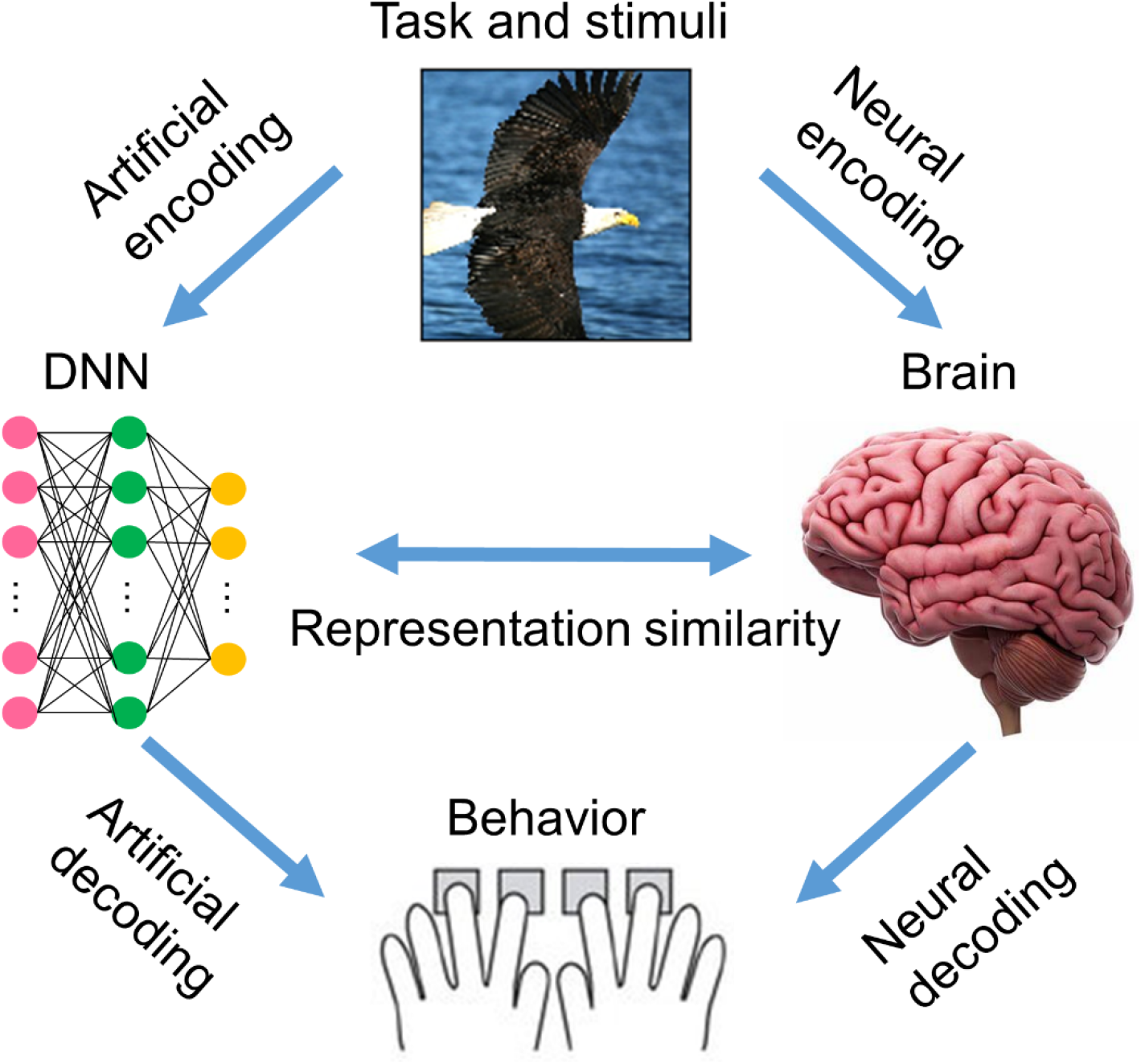
DNNBrain is designed as an integrated toolbox to characterize artificial representations of DNNs and the neural representations of the brain. After stimuli are submitted to both DNNs and human, the artificial neural activity and the biological neural activity can be acquired. By assembling stimuli, artificial activity data, and biological neural activity data together with custom-designed auxiliary IO files, DNNBrain allows users to easily characterize, compare, and visualize representations of DNNs and brains.

- Versatility: DNNBrain supports a diverse range of applications in utilizing DNNs to understand the brain such as accessing DNN representations, mapping DNN representations to the brain, representational similarity analysis (RSA) between DNNs and the brain, and utilizing cognitive neuroscience methods to unveiling DNNs’ black box such as visualizing DNN representations and evaluating behavioral relevance of the representations.
- Usability: DNNBrain provides a command line interface (CLI) and an application programming interface (API) to process DNN and brain imaging data. At the application level, users can conveniently run CLI to conduct typical representation analysis on data. At the programming level, all algorithms and computational pipelines are encapsulated into objects with a high-level interface in the experimental design and data analysis language of neuroscientists. Users can write their own scripts to develop a more flexible and customized pipeline.
- Transparent input/output (IO): DNNBrain supports diverse neuroimaging data formats and multiple meta-data file formats, and can automatically complete data reading, conversions, and writing. Therefore, DNNBrain spares users from specific knowledge about different data formats.
- Open source: DNNBrain is freely available in source and binary forms. Users can access every detail of the DNNBrain implementation, which improves the reproducibility of experimental results, leads to efficient debugging, and gives rise to accelerated scientific progress.
- Portability: DNNBrain, implemented in Python, can run on all major operating systems. It is easy to set up without complicated dependencies on external libraries and packages.

The toolbox is freely available for download^3^ and complemented with an expandable online documentation^4^. As follows, we first introduce the framework of DNNBrain and its building blocks. Then, with a typical application example, we demonstrate the versatility and usability of DNNBrain in characterizing DNNs and in examining the correspondences between DNNs and the brain.

## Overview of the DNNBrain

### The framework of DNNBrain

DNNBrain is a modular Python toolbox that consists of four modules: IO, Base, Model, and Algorithm (Figure 2). The Python language was selected for DNNBrain because it provides an ideal environment for research on DNNs and the brain. First, Python is currently the most commonly-used programming language for scientific computing. A lot of excellent Python libraries have been developed for scientific computing. The libraries used in the DNNBrain are: NumPy for numerical computation^5^, SciPy for general-purpose scientific computing^6^, Scikit-learn for machine learning^7^, and Python imaging library (PIL) for image processing^8^. Second, Python has increasingly been used in the field of brain imaging. Many Python libraries have been developed for brain imaging data analysis such as NiPy^9^ (Millman and Brett, 2007)and fMRIPrep^10^(Esteban et al., 2019). Finally, Python is the most popular language in the field of DL. Python is well supported by the two most popular DNN libraries (i.e., PyTorch^1^ and TensorFlow^2^). Using Python and these DNN libraries, users can build their own DNN in just a couple of lines of code.

**Figure 2.**
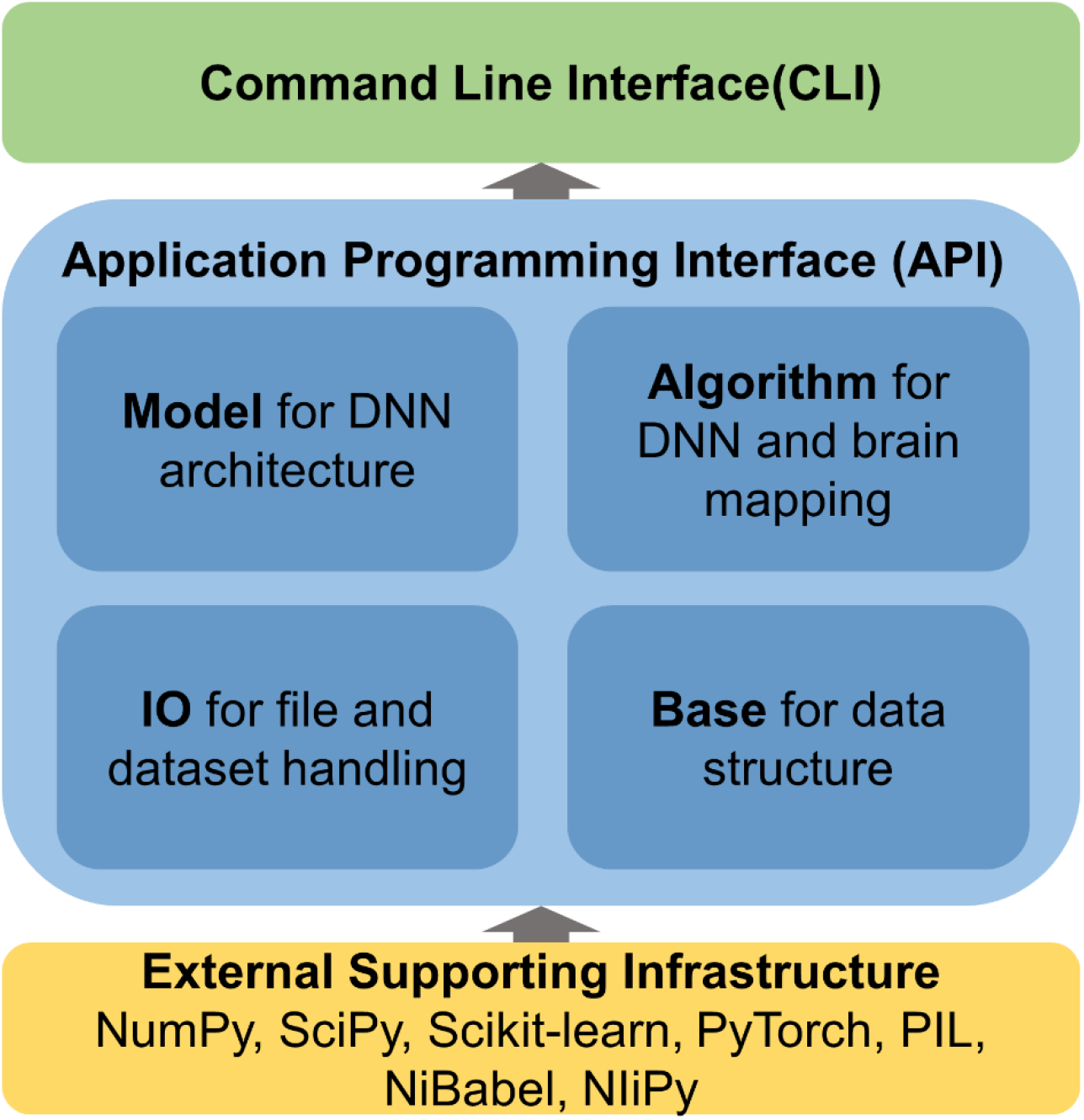
DNNBrain is a modular framework which consists of four modules: IO, Base, Model, and Algorithm. The IO module provides facilities for managing the file-related input and output operations. The Base module defines the base level classes for array computing and data transforming. The Model module holds a variety of DNN models. The Algorithm module defines various algorithms for exploring DNNs and the brain. All modules provide user-friendly APIs. A set of CLIs was developed for a variety of research scenarios.

Supported by a large variety of existing software packages, DNNBrain was designed with a high-level API in the domain language from the cognitive neuroscience. All algorithms and computational pipelines are encapsulated into classes in an object-oriented programming manner. All modules provide user-friendly APIs. Based on the APIs, a set of CLIs was developed for a variety of research scenarios.

### IO module: Organizing datasets in DNNBrain

DNNBrain introduces a few auxiliary file formats to handle various types of scientific data and supporting metadata including stimulus files, DNN mask files, and DNN activation files. With these file formats, users can easily organize their inputs and outputs. The stimulus file is a comma separated values (CSV) text file designed to configure the stimulus information including the stimulus type (image or video), stimulus directory, stimulus ID, stimulus duration, stimulus conditions, and other possible stimulus attributes. The DNN mask file is also a CSV text file designed for users to specify channels and units of interest in analyzing the DNN. Both the stimulus file and the DNN mask file can be easily configured through a text editor. The DNN activation file is a h5py file in which activation values from the specified channels are stored. Besides, DNNBrain uses NiBabel^11^ to access brain imaging files. Almost all common brain imaging file formats, including GIFTI, NIfTI, CIFTI, AFNI, and MGH, are supported.

### Base module: defining the basic data structure

The base module defines the base level objects for data structure and data transformations. Specifically, a set of objects is defined to organize the data from the input stimulus and the output activation data from the DNN, respectively. The data objects were designed as simple as possible while keeping necessary information for further representation analysis. The stimulus object contains stimulus images and associated attributes (e.g., category label), which are read from stimulus files. The activation object holds DNN activation patterns and associated location information (e.g., layer, channel and unit). Beside these data objects, several data transformation objects were also developed, including popular classification and regression models such as generalized linear models, support vector machines, logistic regression and Lasso. All these models were wrapped from the widely used machine learning library scikit-learn^7^.

### Model module: encapsulating DNNs

In DNNBrain, a DNN model is implemented as a neural network model from PyTorch. Each DNN model is a sequential container which holds the DNN architecture (i.e., connection pattern of units) and associated connection weights. The DNN model is equipped with a suit of methods that accesses the attributes of the model and updates the states of the model. PyTorch has become the most popular DL framework because of its simplicity and ease of use in creating and deploying DL applications. At present, several well-known pretrained PyTorch DCNN models^12^ have been adopted into DNNBrain including AlexNet (Krizhevsky et al., 2012), VGG (Simonyan and Zisserman, 2015), GoogLeNet (Szegedy et al., 2014), and ResNet (He et al., 2016) which were pretrained on ImageNet for classification of 1,000 object categorizes^13^.

### Algorithm module: characterizing DNNs and brains

The algorithm module defines various algorithms objects for exploring DNNs. An algorithm object contains a DNN model and corresponding methods to study specific properties of the model. Three types of algorithms are implemented in the DNNBrain. The first one is the gradient descent algorithm for DNN model training wrapped from PyTorch^14^. The second are tools for extracting and summarizing the activation of a DNN model such as principal component analysis (PCA) and clustering. The third type are algorithms to visualize the representations of a DNN including discovering the top stimulus, mapping saliency features of a stimulus, and synthesizing the maximum activation stimulus for a specific DNN channel (Montavon et al., 2018; Nguyen et al., 2019). Each algorithm takes a DNN model and a stimulus object as input which can be imported from a user specified stimulus file.

### The command line interface

At the application level, DNNBrain provides several workflows as a command-line interface, including accessing the DNN representations, visualizing the DNN representations, evaluating the behavioral relevance of the representations, and mapping the DNN representations to the brain. Users can conveniently run command lines to perform typical representation analysis on their data.

## Methods

### DNN model: AlexNet

AlexNet is one of the most influential DCNNs that demonstrated that DCNNs could significantly increase ImageNet classification accuracy by a significant stride for the first time in the 2012 ImageNet challenge (Krizhevsky et al., 2012). AlexNet is composed of five convolutional (Conv) layers and three fully connected (FC) layers that receive inputs from all units in the previous layer (Figure 3A). Each convolutional layer is generally comprised of a convolution, a rectified linear unit function (ReLU), and max pooling operations. These operations are repeatedly applied across the image. In the paper, when we refer to Conv layers, we mean the output after the convolution and ReLU operations.

**Figure 3.**
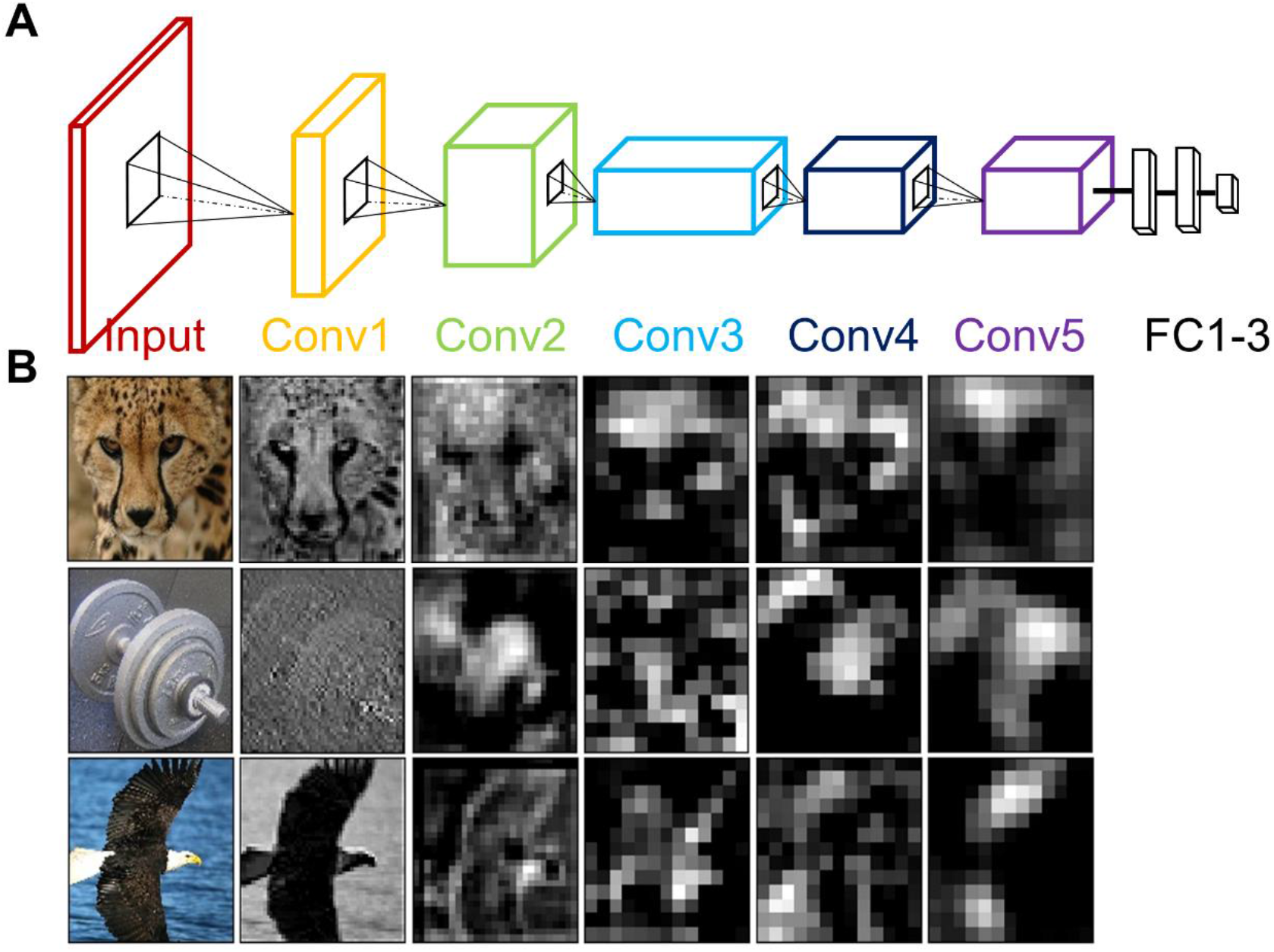
AlexNet architecture and the example unit activity patterns. **(A)** AlexNet consists of five Conv layers followed by three FC layers and finally a 1000-way softmax classifier. **(B)** The activation maps from each five Conv layers of AlexNet were extracted for three example images (cheetah, dumbbell, and bald eagle). The presented channels are those showing the maximal mean activation for that example image within each of the five Conv layers.

### BOLD5000: stimulus and neuroimaging data

The BOLD5000 is a large-scale publicly-available human fMRI dataset in which four participants underwent slow event-related BOLD fMRI while viewing approximate 5,000 distinct images depicting real-world scenes (Chang et al., 2019). The stimulus images were drawn from three most commonly-used computer vision datasets: 1,000 hand-curated indoor and outdoor scene images from the Scene UNderstanding dataset (Xiao et al., 2010), 2,000 images of multiple objects from the Common Objects in Context dataset (Lin et al., 2015), and 1,916 images of mostly singular objects from the ImageNet dataset (Deng et al., 2009). Each image was presented for 1 s followed by a 9 s fixation cross. Functional MRI data were collected using a T2*-weighted gradient recalled echo planar imaging multi-band pulse sequence (In-plane resolution = 2 × 2 mm; 106 × 106 matrix size; 2 mm slice thickness, no gap; TR = 2000 ms; TE = 30 ms; flip angle = 79 degrees). The scale, diversity and naturalness of the stimuli, combined with a slow event-related fMRI design, make BOLD5000 an ideal dataset to explore DNNs and brain representations of a wide range of visual features and object categories. The raw fMRI data were preprocessed with the fMRIPrep pipeline including motion correction, linear detrending, and spatial registration to the native cortical surface via boundary-based registration (Esteban et al., 2019). No additional spatial or temporal filtering was applied. For a complete description of the experimental design, fMRI acquisition, and the preprocessing pipeline, see (Chang et al., 2019).

The BOLD response maps for each image were firstly estimated from the preprocessed individual fMRI data through a general linear model in the subject-native space and then transformed into fsLR space using ciftify (Dickie et al., 2019). The response maps of each image were finally averaged across four subjects in the fsLR space and used for further analyses.

## Results

We demonstrated the functionality of DNNBrain on AlexNet and the BOLD5000 dataset. Specifically, we accessed the DNN activation of the images from BOLD5000, probed the category information represented in each DNN layer, mapped the DNN representations onto the brain, and visualized the DNN representations. These examples do not aim to illustrate the full functionalities available from DNNBrain, but rather to sketch out how DNNBrain can be easily used to examine DNNs’ and brain’s representations in a realistic study. All the analyses were implemented in both API and CLI levels. The code can be found in the DNNBrain online documentation^4^.

### Scanning DNNs

To examine the artificial representations of the DNN, we first scanned the DNN and to obtain its neural activities, just as we scan the human brain using brain imaging equipment. DNNBrain provides both API and CLI to extract the activation states for user-specified channels of a DNN. Figure 3 shows the activation patterns of three example images (cheetah, dumbbell, and bald eagle) from the channels of AlexNet which showed the maximal mean activation within each of the five Conv layers, revealing that the DNN representation of the image became more abstract along the depth of the layers.

### Revealing information presented in DNN layers

To reveal whether specific stimuli attributes or behavioral performances are explicitly encoded in a certain layer of a DNN, a direct approach is to measure to what degree is the representation from the layer useful for decoding them. Linear decoding models (classifier or regression) were implemented in DNNBrain to fulfill this. Here, we manually sorted the BOLD5000 stimulus images into binary categories (animate versus inanimate) according to salient objects located in each image, and examined how animate information are explicitly encoded in AlexNet. In total, 2,547 images were labeled as animate, and 2,369 inanimate. We trained a logistic regression model on the artificial representation from each Conv layer of AlexNet to decode the stimulus category. To avoid the risk of overfitting with limited training data, the dimension (i.e. the number of units) of the activation pattern from each layer was reduced by PCA to retain the top 100 components. The accuracy of the model was evaluated with a 10-fold cross validation. As shown in Figure 4, the classification accuracy progressed with the depth of Conv layers, indicating higher layers encode more animate information than lower layers. Moreover, the ReLU operation within each convolutional layer plays a significant role in improving the representation capacities for animate information.

**Figure 4.**
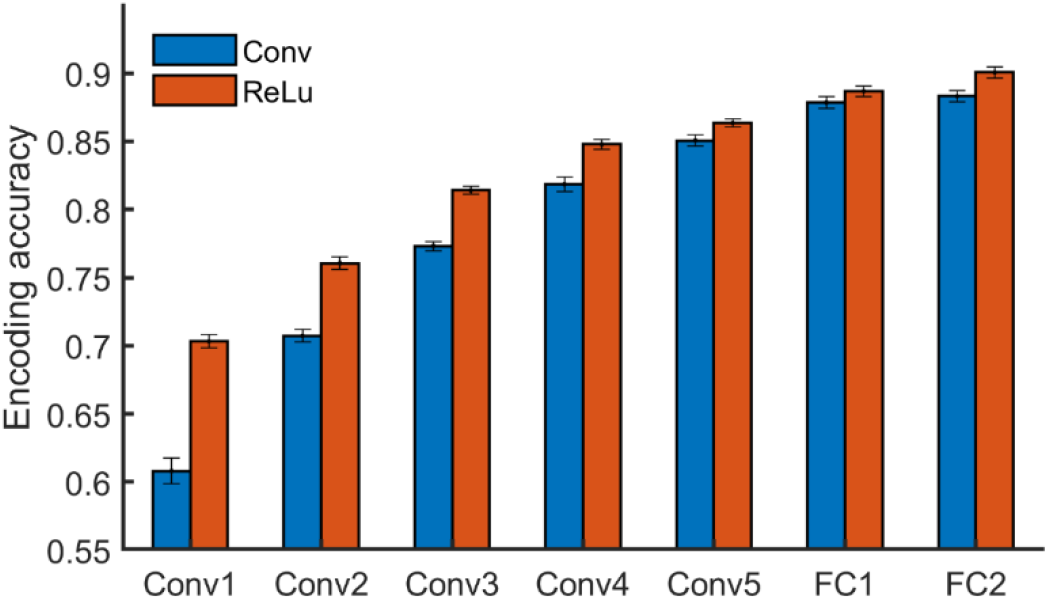
DNNBrain probes (or decodes) the explicit representation contents of layers of interest in a DNN using linear models. On BOLD5000 stimuli, a logistic regression model revealed that the higher a layer is, the more animate information it encoded.

### Mapping representations between a DNN and the brain

A growing body of studies is testing DNNs as a model of brain information processing. For example, several recent studies mapped the artificial representations from a DNN optimized for object classification task to the primate ventral visual stream and revealed that internal representations of DNNs provide the best current models of representations of visual images in the inferior temporal cortex in humans and monkeys (Lindsay, 2020). DNNBrain supports two kinds of analyses to link DNN artificial representations to brain representations: encoding model (EM) (Naselaris et al., 2011) and RSA (Kriegeskorte et al., 2008).

The EM aims to find linear combinations of DNN units to predict the response of a neuron or voxel in the brain. The linear model is preferred because how the two kinds of representations are similar in an explicit format (i.e. a linear transform) is primarily concerned for researchers (Yamins et al., 2014; Wen et al., 2018). Here, we used voxel-wise EM to check how the human ventral temporal cortex (VTC) encodes the representations from the Conv layers of AlexNet. The VTC region was defined by merging the areas V8, FFC, PIT, VVC, and VMV from HCP MMP 1.0 (Glasser et al., 2016). For each voxel within the VTC, five EMs were constructed using the artificial representation from each of five Conv layers in AlexNet. The encoding accuracy was evaluated with the Pearson correlation between the measured responses and the predicted responses from the EM using a 10-fold cross validation procedure on BOLD5000 dataset. Two main findings were revealed (Figure 5A). Firstly, the overall encoding accuracy of the VTC gradually increased for the hierarchical layers of AlexNet, indicating that as the complexity of the visual representations increase along the DNN hierarchy, the representations become increasingly VTC-like. Second, the encoding accuracy varied greatly across voxels within the VTC for the artificial representations of each AlexNet layer, indicating the VTC may organize in distinct functional modules, each preferring different kinds of features.

**Figure 5.**
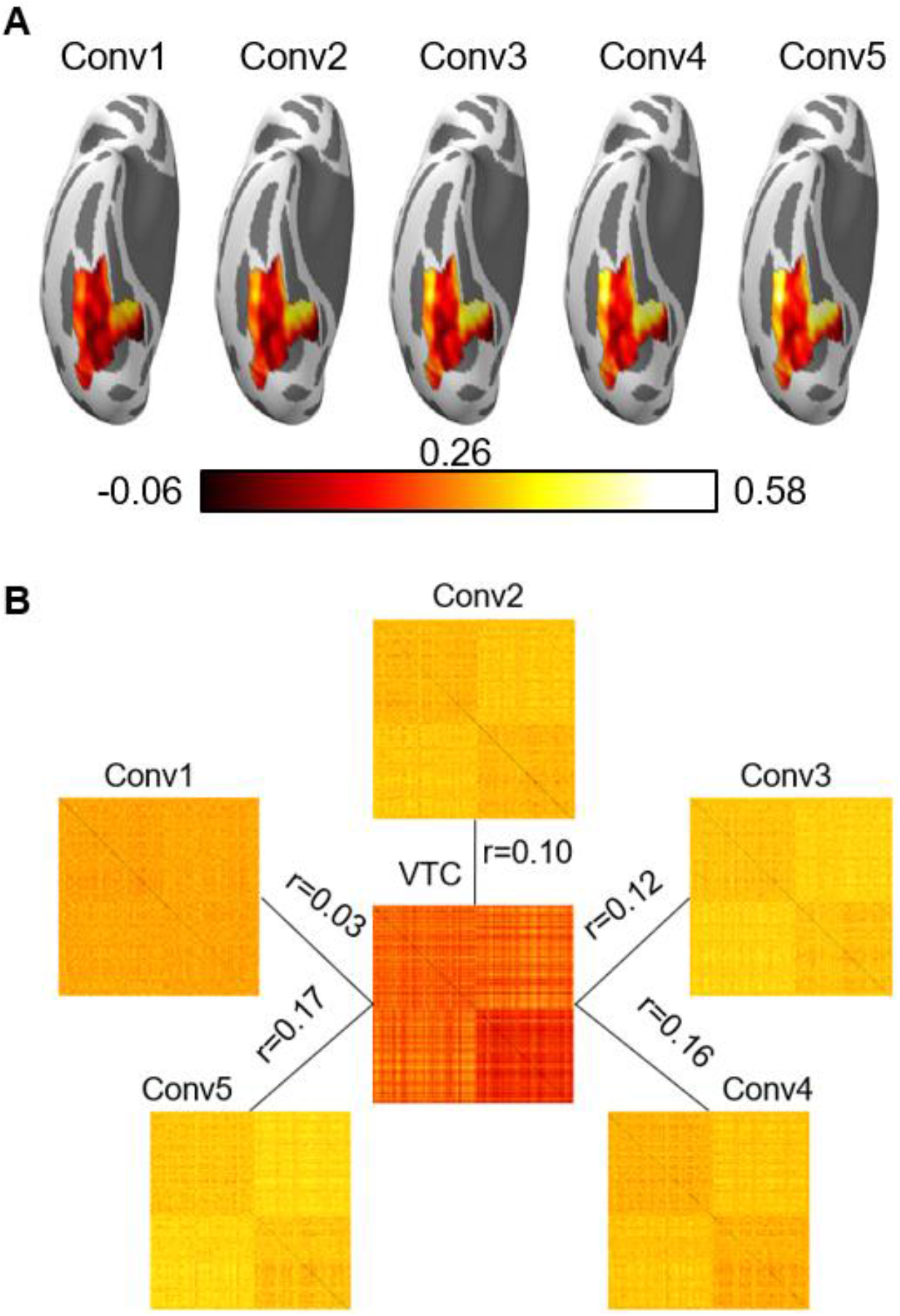
Both the encoding model and the representational similarity analysis are implemented in DNNBrain to help researchers to examine the correspondence between the DNN and brain representations. **(A)** The encoding accuracy map from voxel-wise encoding models of predicting the VTC BOLD responses using the artificial representation from the Conv layers of AlexNet. **(B)** The RDMs for the BOLD5000 stimuli computed on the artificial representation from Conv layers of AlexNet and brain activation patterns from the human VTC. The representation distance between each pair of images was quantified as the correlation distance between their representation. The representation similarity between DNN and brain is further calculated as the Pearson correlation between their RDMs.

Instead of predicting brain responses directly, RSA compares the representations of the DNN and that of the brain using a representational dissimilarity matrix (RDM) as a bridge: the RDMs are first created to measure how similar the response patterns are for every pair of stimuli or conditions using the multivariate response patterns from the DNN and the brain, respectively. The representation similarity between the DNN and the brain is further calculated as the correlation between their RDMs (Khaligh-Razavi and Kriegeskorte, 2014; Cichy et al., 2016). As an example, the RDMs of the BOLD5000 stimuli are shown in Figure 5B, which were calculated on the BOLD representation from the VTC and the artificial representation from each of the five Conv layers of AlexNet. First, the RDM from AlexNet revealed that the category representations gradually emerge along the hierarchy. Second, the artificial representations of AlexNet are increasingly resembling the neural representation from the human VTC.

### Visualizing features from DNNs

DNNs are a kind of complex non-linear transformation that does not provide any explicit explanation of their internal workings. Identifying relevant features contributing the most to the responses of an artificial neuron is central to understand what exactly each neuron has learned (Montavon et al., 2018; Nguyen et al., 2019). In DNNBrain, three approaches were implemented to help users examine the stimulus features that an artificial neuron prefers. The first approach is top stimulus discovering that finds the top images with the highest activations for a specific neuron (or unit) from a large image collection (Zeiler and Fergus, 2014; Yosinski et al., 2015). The second approach is saliency mapping that computes gradients on the input images relative to the target unit by a backpropagation algorithm. It highlights pixels of the image that increase the unit’s activation most when its value changes (Simonyan et al., 2014; Springenberg et al., 2015). The third approach is optimal stimulus synthesizing which synthesizes the visual stimulus from scratch guided by increasing activation of the target neuron (Erhan et al., 2009; Nguyen et al., 2016). It offers advantages over the top stimulus discovering and saliency mapping because it avoids the risks that effective images that could activate the neuron may not exist in the stimulus set.

We used DNNBrain to visualize the preferred features for three output units of AlexNet (i.e., ostrich, peacock, and flamingo) as an example. The output units were selected as examples because the produced features for them are easy to check (i.e., each unit corresponds to a unique category). These procedures essentially work for any unit in a DNN. As shown in Figure 6A, the top stimulus was correctly found from 4916 BOLD5000 images for each of three units: every top stimulus contains the object in the correct category. The saliency maps highlight the pixels in the top stimuli that contribute to the activation of the neurons most (Figure 6B). Finally, the images synthesized from scratch correctly emerge objects of the corresponding category (Figure 6C). In summary, these three approaches are able to reveal the visual patterns that a neuron has learned on various levels and thus provide a qualitative guide to neural interpretations.

**Figure 6.**
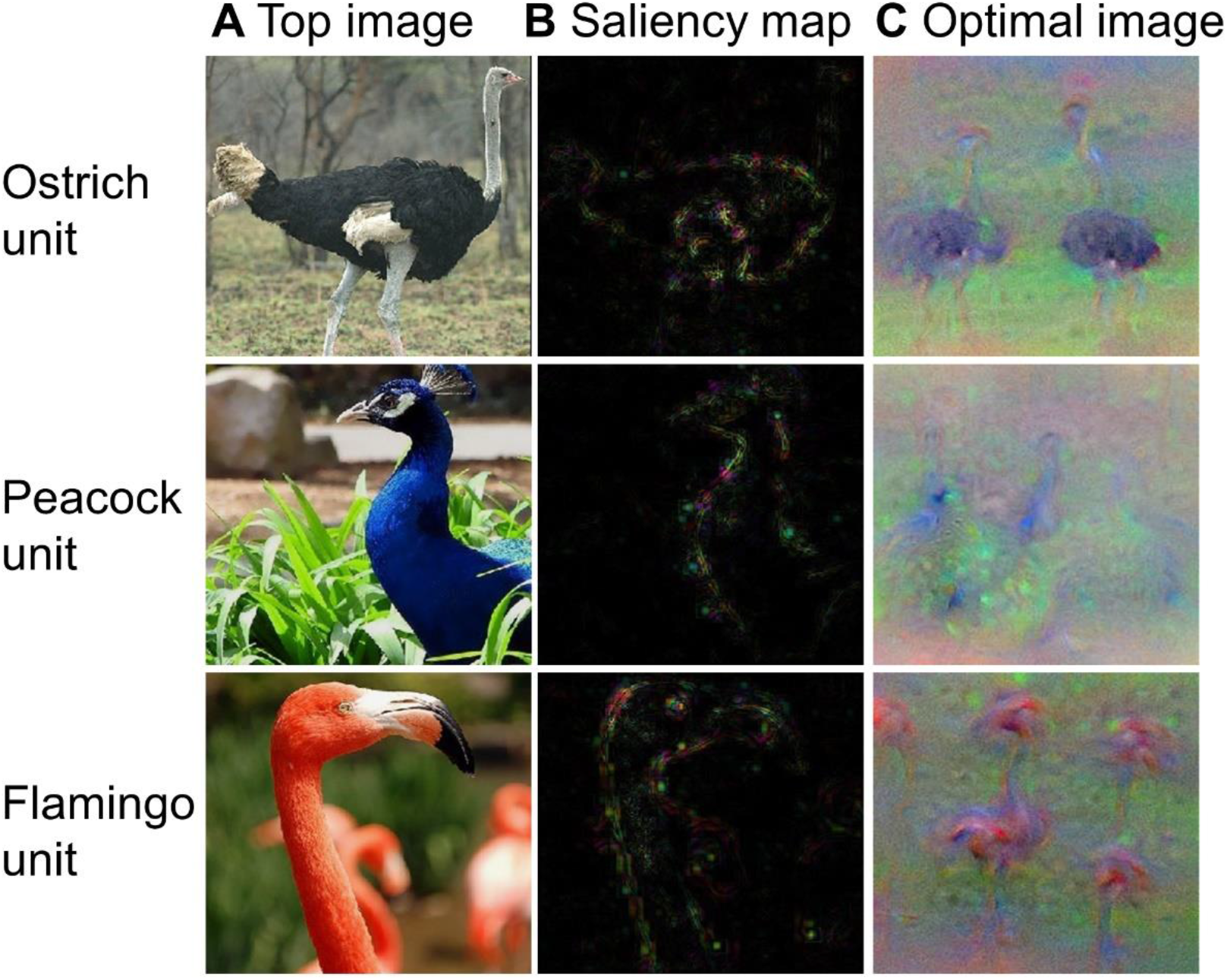
The top stimuli, saliency maps and synthesized images for three output units of AlexNet. **(A)** The top stimuli discovered from the BOLD5000 dataset. **(B)** The saliency maps computed for the top stimuli presented in (A). **(C)** The images synthesized from scratch guided by increasing the activation of corresponding neurons.

### Other analyses DNNBrain supported

Besides the functionality illustrated in the above examples, DNNBrain also provides many other flexible pipelines for neuroscience-orientated analysis of DNNs including ablation analysis of individual units and estimation of the empirical receptive field of a unit (Zhou et al., 2014). It also comes with a variety of utilities such as image processing tools used for converting image data types between PyTorch tensor, NumPy array, and PIL image objects, translating and cropping images, etc. A set of utilities that helps users training a new model or doing transfer learning was also provided. Visit the DNNBrain documentation page for details^4^.

## Discussion

DNNBrain integrates well-established DNN software and brain imaging packages to enable researchers to conveniently map the representations of DNNs and brains, and examining their correspondences (Figure 1). DNN models provide a biologically plausible account of biological neural systems, and show great potential for novel insights into the neural mechanisms of the brain. On the other hand, experimental paradigms from cognitive neuroscience provide powerful approaches to pry open the DNNs’ black box, which promotes the development of explainable DNNs. DNNBrain as a toolbox that is explicitly tailored toward integrated mapping between DNNs and the brain will likely accelerate the merge of AI and neuroscience.

DNNBrain integrates many of the currently most popular pretrained DCNN models. With the advance of the interplay of neuroscience and DNN communities, new DNN models are constantly emerging and will be included into DNNBrain in the future. For example, a generative adversarial network could be introduced into DNNBrain to help users to reconstruct external stimuli (Shen et al., 2019; VanRullen and Reddy, 2019) and to synthesize preferred images for neurons or brain areas (Ponce et al., 2019) according to their dynamic brain activities. Besides, there are a few issues that we would like to address in the future. First, DNNBrain until now only supports DNN models from PyTorch, which limits users to study DNNs constructed under other frameworks within DNNBrain. We next will make great efforts to integrate the TensorFlow framework into DNNBrain. Second, only fMRI data is currently well supported in DNNBrain because the organization of other types of brain imaging data have not yet been well standardized (Gorgolewski et al., 2016). As the standardization of electrophysiology data progresses (Niso et al., 2018; Pernet et al., 2019), we would very much like to extend DNNBrain to support magnetoencephalography, electroencephalography, multiunit recordings, and local field potentials. Finally, DNNBrain mainly supports the exploration of pretrained models trained on external stimuli. A recent advance demonstrated the brain representation can provide additional and efficient constraints on DNN constructions (McClure and Kriegeskorte, 2016). Therefore, it would be a good attempt to equip DNNBrain with tools in the future to fuse brain activities and external tasks/stimuli to create DNN models that are more closely resembling the human brain.

## Code and data availability

Our toolbox is freely available via github^3^. The code used in this tutorial and additional documentation is available for via readthedocs^4^. All data are freely provided by the BOLD5000 Project^15^ and available from Kilthub^16^ or OpenNeuro^17^.

## Footnotes

1 https://pytorch.org

2 https://www.tensorflow.org

3 http://github.com/BNUCNL/dnnbrain

4 http://dnnbrain.readthedocs.io

5 https://numpy.org

6 https://www.scipy.org

7 https://scikit-learn.org

8 http://pythonware.com/products/pil

9 https://nipy.org

10 https://fmriprep.org

11 https://nipy.org/nibabel

12 https://github.com/pytorch/vision

13 http://image-net.org

14 https://pytorch.org/docs/stable/optim.html

15 https://bold5000.github.io

16 https://figshare.com/articles/BOLD5000/6459449

17 https://openneuro.org/datasets/ds001499/versions/1.3.0

## Notes

**Conflict of interest:** The authors declare that the research was conducted in the absence of any commercial or financial relationships that could be construed as a potential conflict of interest.

### Competing Interest Statement

The authors have declared no competing interest.

